# Titin cleavage in living cardiomyocytes induces sarcomere disassembly but does not trigger cell proliferation

**DOI:** 10.1101/2025.04.22.645658

**Authors:** Maria Rosaria Pricolo, Miguel A. López-Unzu, Natalia Vicente, Cristina Morales-López, Carla Huerta-López, Wendy Pérez-Franco, Andra C. Dumitru, Francisco M. Espinosa, Mateo I. Sanchez, Ricardo Garcia, Roberto Silva-Rojas, Elías Herrero-Galán, Jorge Alegre-Cebollada

## Abstract

**Aims:** Adult mammalian hearts have limited regenerative capacity due to the inability of cardiomyocytes to proliferate, a major clinical hurdle in contemporary cardiology. The presence of highly organized, contractile sarcomeres has long been considered an impediment for cardiomyocyte division. Indeed, sarcomere disassembly is a crucial step to complete the cell cycle in the few situations where cardiomyocytes have been observed to proliferate. However, whether sarcomere disassembly can per se trigger cell cycle re-entry remains unknown, a possibility that we have tested here.

**Methods and results:** We have engineered a system to induce sarcomere disassembly in living murine cardiomyocytes based on the specific cleavage of the structural protein titin by tobacco etch virus protease (TEVp). Although isolated neonatal cardiomyocytes with disassembled sarcomeres remain viable and retain low-amplitude contractile activity, our results show no evidence of increased cardiomyocyte proliferation in targeted cells, as indicated by analyses of markers of DNA synthesis and cytokinesis. We obtain equivalent results when titin is cleaved in the adult myocardium *in vivo*.

**Conclusion:** The removal of sarcomere structural barriers is necessary, but not sufficient, for cardiomyocyte proliferation, which implies that additional factors are required for cardiomyocytes to undergo cell division.

**Translational perspective:** There is a clinical need to identify therapeutic strategies that promote cardiac regeneration through the proliferation of cardiomyocytes that survive an injury to the heart, for instance after myocardial infarction. Based on the observation that cardiomyocytes require sarcomere disassembly for proliferation, we have examined if the sole disassembly of sarcomeres is enough to promote cell division in cardiomyocytes. Our work demonstrates a strategy to induce specific sarcomere disassembly, which, however does not result in increased proliferative capacity of cardiomyocytes. These results imply that additional factors need to be considered to promote cardiomyocyte proliferation by facilitating sarcomere disassembly.

## Introduction

Heart failure, a leading cause of morbidity and mortality worldwide, is mostly the long-term consequence of conditions causing massive and irreversible loss of cardiomyocytes, such as myocardial infarction^1^. To improve clinical outcomes in these conditions, extensive efforts are being directed to promote cardiac regeneration through the induction of proliferation of surviving cardiomyocytes^2^. However, this endeavor faces significant challenges. Whereas mammalian cardiomyocytes do proliferate during embryonic and fetal development, the vast majority of them exit the cell cycle within the first few days/weeks after birth and remain persistently refractory to proliferation from that moment onwards ^3–5^. Indeed, the natural response of adult mammalian hearts to cardiomyocyte loss is to replace necrotic tissue by a fibrotic scar and to induce hypertrophic growth of the remaining cardiomyocytes, which however typically fail to restore global contractile function^6,7^.

Different from mammals, the heart from a number of vertebrates including zebrafish, newts, and axolotls, can regenerate completely after injury via the activation of cardiomyocyte proliferation^8–10^. Based on these observations and on evidence that a limited fraction of mammalian cardiomyocytes can re-enter the cell cycle to some degree after myocardial injury^11,12^, several groups have identified a few pathways and molecules that induce proliferation of adult murine cardiomyocytes correlating with improved cardiac function following myocardial infarction ^13–18^. Disassembly of sarcomeres, the contractile units of terminally differentiated cardiomyocytes^19^, is a feature common to both natural and induced cardiomyocyte proliferation^20^ (**Figure 1A**). Importantly, sarcomere disassembly is required for cardiomyocytes to proliferate, since inhibition of actin-capping adducin proteins or MMP-2 protease, two mediators of sarcomere disassembly, blunts cardiomyocyte proliferation^21,22^. These results support the traditional view that the dense arrays of sarcomeres in cardiomyocytes (i.e. the myofibrils) are an intrinsic barrier to cell division, potentially involving crosstalk with other cytoskeletal components and mitogenic signaling pathways^20^.

**Figure 1.**
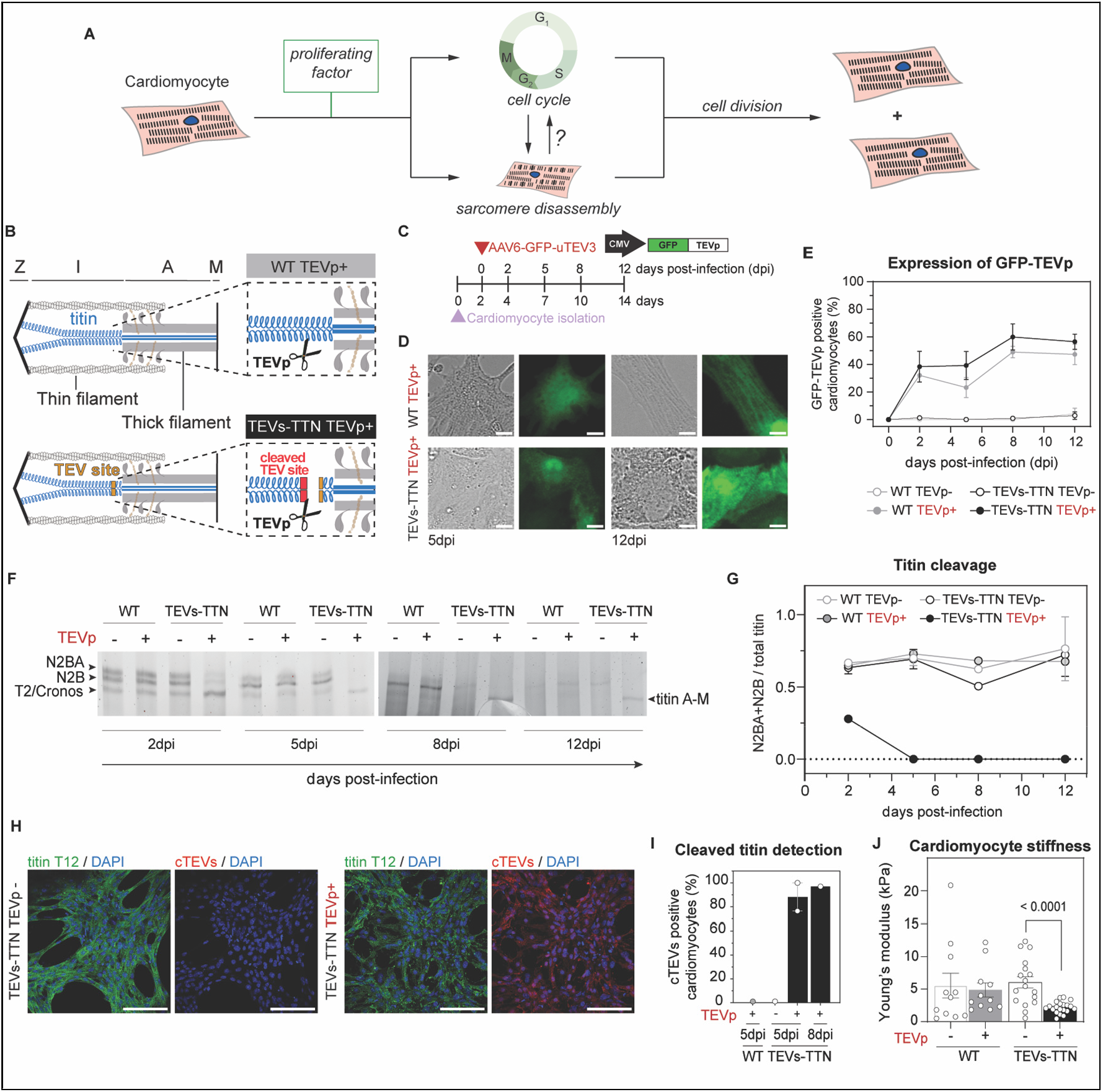
TEVp-induced cleavage of titin in living cardiomyocytes. **(A)** Scheme of cardiomyocyte proliferation, which requires cell cycle progression and sarcomere disassembly. This work examines whether sarcomere disassembly is enough to trigger progression in cell cycle. **(B)** Schematic representation of the titin cleavage strategy. TEVp recognizes the knocked-in TEVs within the I-band of titin in TEVs-TTN cardiomyocytes and cleaves titin. In contrast, titin remains uncleaved in WT cells. **(C)** Experimental design. Upon isolation, neonatal mouse cardiomyocytes are infected *in vitro* with GFP-TEVp-expressing AAV6 particles and analyzed at different days post infection. **(D)** Bright-field (*left*) and epifluorescence (*right*) representative images of living neonatal cardiomyocytes. Scale bars, 10 µm. **(E)** TEVp expression quantification from native GFP fluorescence (WT N=3-4; TEVs-TTN N=4-8). **(F)** 1.8% polyacrylamide SDS-PAGE electrophoresis to analyze titin content. **(G)** Quantification of the proportion of the T2, Cronos and A-M titin bands with respect to all titin bands (N=2-3). **(H)** Detection and (**I**) quantification of cleaved titin with anti cTEVs. Scale bars, 100 µm. **(J)** Stiffness of skinned cardiomyocytes measured using AFM. Each dot represents the mean value of several measurement for a single cell (N=2 isolations of cardiomyocytes).

Here, we hypothesized that the sole disassembly of sarcomeres could be enough to promote cardiomyocyte proliferation (**Figure 1A**). In support of this hypothesis, cardiomyocytes devoid of sarcomeres are more proliferative^23^, soft extracellular matrices disorganize sarcomeres and induce cardiomyocyte cell-cycle re-entry^24^ and overexpression of adducin proteins can trigger sarcomere disassembly in cardiomyocytes correlating with increased levels of proliferation markers^22^. Based on data that severing titin, a central structural component of myofibrils^25,26^ (**Figure 1B**), destabilizes skeletal muscle sarcomeres^27,28^, we set out to investigate whether titin cleavage could be exploited to elicit sarcomere disassembly and subsequent proliferation of targeted cardiomyocytes. Our results demonstrate that full titin cleavage does indeed cause sarcomere disassembly in otherwise viable cells; however, contrary to our motivating hypothesis, we find no evidence of increased proliferation of cardiomyocytes containing cleaved titin.

## Results

### Induction of specific titin cleavage in living cardiomyocytes

For our experiments, we exploited a homozygous knock-in mouse model that contains a recognition site for tobacco etch virus protease (TEVp) in the elastic I-band region of titin (TEVs-TTN mice)^29^ (**Figure 1B**). The ability of TEVp to cleave cardiac titin from TEVs-TTN sarcomeres has been demonstrated adding purified TEVp to demembranated preparations^27,29,30^. Here, to implement titin cleavage in living cells, we first infected neonatal TEVs-TTN cardiomyocytes with AAV6 viral particles expressing different forms of TEVp under the control of the constitutive CMV promoter. Among the four different TEVp constructs tested, we find optimal expression and cytosolic localization of GFP-uTEV3 (hereafter referred to as GFP-TEVp), which contains an improved version of TEVp obtained by directed evolution^31^ (**Figure S1A**). We used GFP-TEVp-expressing AAV6 particles for all subsequent experiments.

We tracked expression of GFP-TEVp in both wild-type (WT) and TEVs-TTN neonatal cardiomyocytes at different days post infection (dpi). In both genotypes, GFP positive cells are already evident at 2 dpi reaching ∼60% of cardiomyocytes from 8 dpi (**Figures 1C-E, S1B**). Next, we examined the levels of titin cleavage by 1.8% polyacrylamide SDS-PAGE electrophoresis. As expected, in all WT and untreated TEVs-TTN control cells, we detect three major titin bands corresponding to full-length N2BA and N2B isoforms and the higher electrophoretic-mobility band originating from the Cronos isoform and/or the T2 degradation fragment^26^ (**Figure 1F**). In contrast, AAV-infected TEVs-TTN samples show just one major titin band of electrophoretic mobility compatible with the C-terminal titin fragment derived from TEVp activity^29^ (A-M titin, similar mobility to Cronos/T2, **Figure 1F**). Quantification demonstrates that full-length titin is completely absent in TEVs-TTN samples from 5 dpi (**Figure 1G**). In agreement with this result, from this time point the vast majority of TEVs-TTN cardiomyocytes (identified as cells showing titin staining with T12 antibody^32^) are also positive for anti-cleaved TEVs (cTEVs) antibody^28^ **(Figure 1H**,**I**), which highlights limited sensitivity of GFP-TEVp detection by epifluorescence (**Figure 1E**). Finally, considering that titin cleavage with purified TEVp results in lower cardiomyocyte stiffness in demembranated preparations^27^, we used atomic force microscopy (AFM) to determine whether cell softening also occurs when cleavage is induced in living cells. Results show that the Young’s modulus of demembranated cardiomyocytes carrying cleaved titin is 2.2 ± 0.8 KPa, around three times lower than the three control groups (**Figure 1J**). These data are in line with extensive evidence collected from *ex vivo* systems supporting that titin is a major contributor to the passive stiffness of cardiomyocytes^25,30^.

In combination, our results demonstrate successful implementation of titin cleavage in living cardiomyocytes. Severing of titin is complete even at low GFP-TEVp levels that cannot be detected by conventional epifluorescence, illustrating the catalytic nature of our protease-based approach. Importantly, we find no effects of GFP-TEVp expression in cardiomyocytes that are devoid of the TEVp recognition site in titin, which supports the specificity of the phenotypes we detect upon GFP-TEVp expression in TEVs-TTN cardiomyocytes.

### Sarcomere disassembly upon titin cleavage in otherwise viable cardiomyocytes

Having verified the full cleavage of titin in AAV-infected TEVs-TTN cardiomyocytes, we next examined the structural integrity of targeted sarcomeres. Z-disk staining with T12 anti-titin antibody indicates loss of normal sarcomere pattern in AAV-infected TEVs-TTN cardiomyocytes, but not in any of the three control groups (**Figures 2A,B**, **S2A**). We quantified sarcomere disassembly both using an automatic analysis of the anisotropy of T12 fluorescence signal (**Figures 2C, S2B**), and by blinded inspection of fluorescence images to classify structural integrity of sarcomeres (**Figure 2D**). Both analyses readily capture progressive sarcomere disorganization in AAV-infected TEVs-TTN cardiomyocytes. Indeed, most TEVs-TTN cardiomyocytes in the AAV-infected group show at least partially disassembled sarcomeres already at 2 dpi; no intact sarcomeres are detected in this experimental group at 8 dpi (**Figure 2D**).

**Figure 2.**
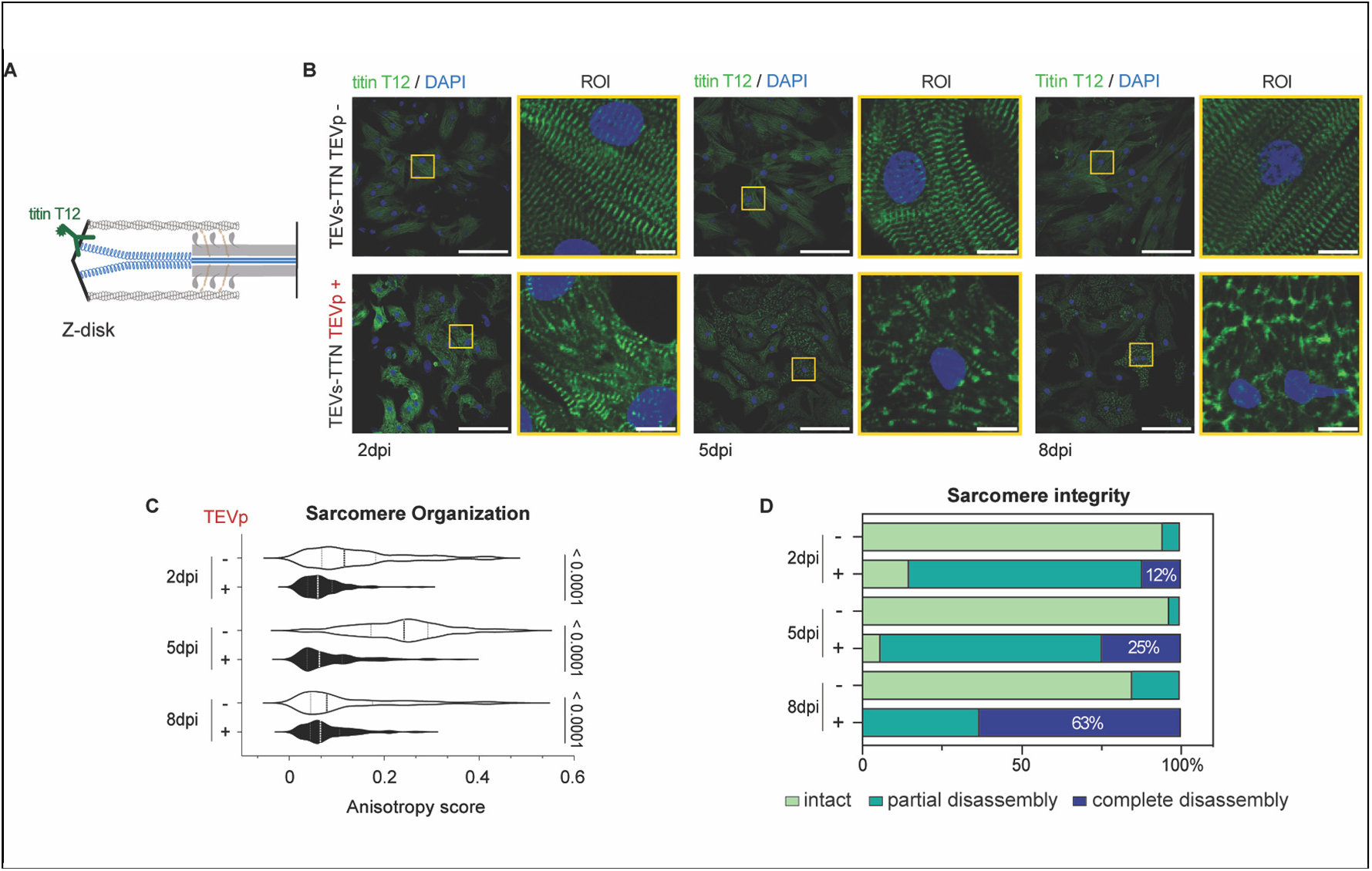
Sarcomere disassembly upon titin cleavage. **(A)** The T12 titin epitope localizes to the Z-disk of sarcomeres. **(B)** Representative immunofluorescence images of TEVs-TTN cardiomyocytes expressing or not GFP-TEVp. Scale bars, 100 µm. ROI scale bars, 10 µm. **(C)**. Sarcomere organization quantified as anisotropy of T12 signal in immunofluorescences (N=2-3). **(D)** Qualitative estimation of sarcomere integrity (N=2).

We extended our analysis to other relevant components of the sarcomere (**Figure 3A**). Similar to T12 staining, Z-disk scaffold *α*-actinin shows a disorganized, non-sarcomeric pattern resembling disconnected Z-bodies already at 2 dpi^21^ (**Figure 3B, S2C**). Staining of troponin T, a thin filament component, is also devoid of striated pattern in AAV-infected TEVs-TTN cardiomyocytes, although in this case remnant fibrillar structures are seen even at 8 dpi (**Figure 3B, S2C**). Finally, targeted cardiomyocytes show diffuse staining of the TTN-9 M-band titin epitope (**Figure 3C**).

**Figure 3.**
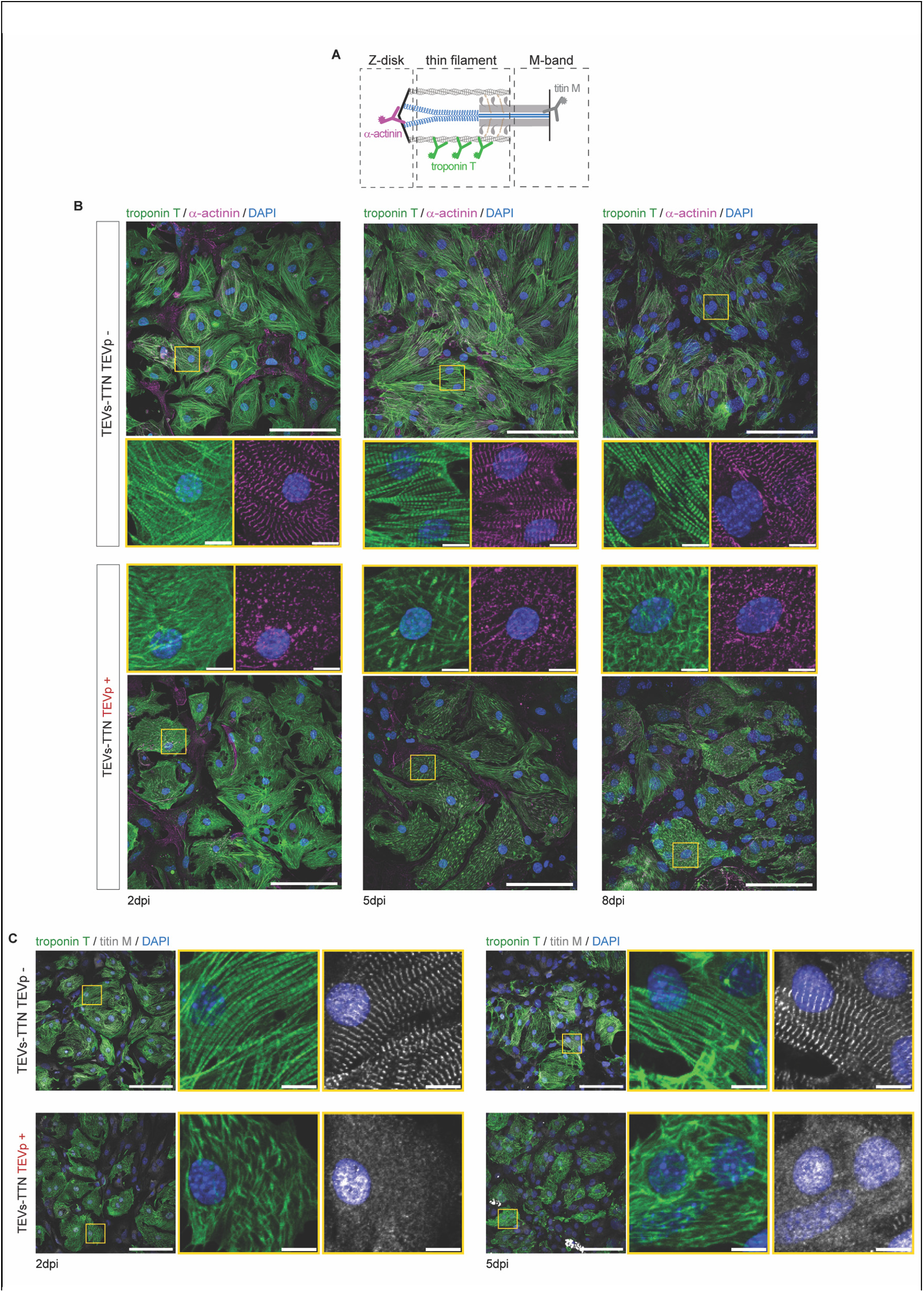
Thin filament and M-line disassembly upon titin cleavage. **(A)** Sarcomere epitopes used in immunofluorescence imaging to investigate sarcomere disarray. **(B**,**C)** Representative immunofluorescence images of TEVs-TTN cardiomyocytes expressing or not GFP-TEVp. Scale bars, 100 µm. ROI scale bars, 10 µm.

We asked ourselves whether sarcomere disassembly upon titin cleavage affects cardiomyocyte viability. Using the 3-[4,5-dimethylthiazol-2-yl]-2,5 diphenyl tetrazolium bromide (MTT) assay, we find no evidence of altered metabolic activity in TEVs-TTN cardiomyocytes at 5 dpi (**Figure 4A**). Similarly, we detect no indication of apoptosis using terminal deoxynucleotidyl transferase dUTP nick end labelling (TUNEL) assays at 5 dpi (**Figure 4B**). Furthermore, tracking spontaneous cardiomyocyte contractility by light microscopy (**Figure 4C,D**), we observe that most AAV-infected TEVs-TTN cardiomyocytes are able to beat, although with much reduced amplitude as expected from the observed loss of sarcomeres (**Figure 4E**,**F, Supplementary Videos S1**,**S2**). This effect is accompanied by lower duration of contractions at longer time points (**Figure 4G**).

**Figure 4.**
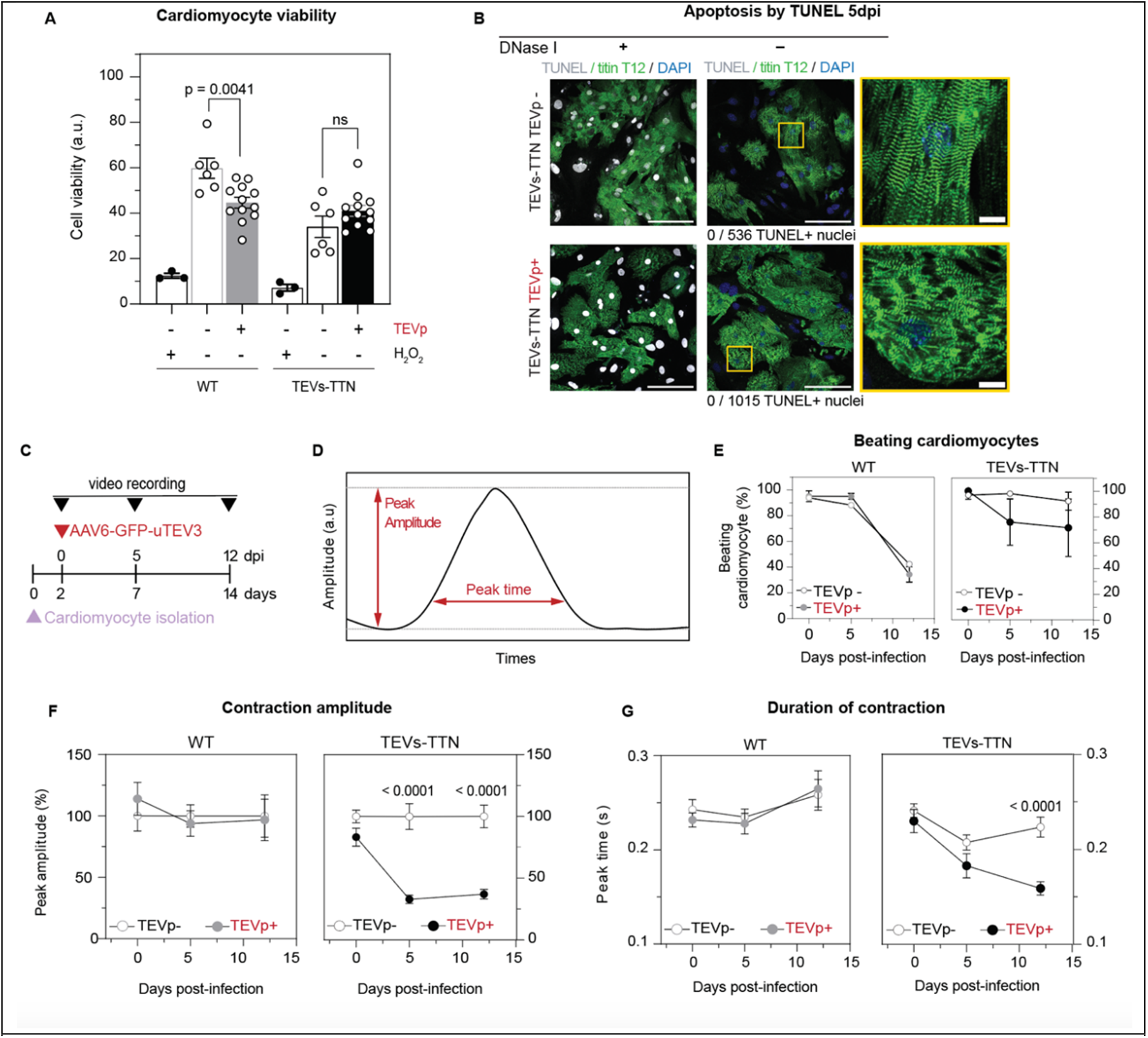
Beating alteration in GFP-TEVp-expressing TEVs-TTN cardiomyocytes in the absence of cell death. **(A)** Cardiomyocyte viability determined via the MTT assay at 5 dpi. Cells incubated with hydrogen peroxide (H_2_O_2_) are used as positive control. **(B)** Representative images of TEVs-TTN cardiomyocyte apoptosis detected by TUNEL. Positive control samples are incubated with DNase I. Scale bars, 100 µm. ROI scale bar, 10 µm. **(C)** Schematic of experiment to monitor cardiomyocyte contraction. 0 dpi samples were measured before infection. **(D)** Peak amplitude and time are analyzed with MYOCYTER^51^. **(E)** Fraction of spontaneously beating WT (*left*) and TEVs-TTN (*right*) cardiomyocytes, expressing or not GFP-TEVp (N=3). **(F)** Contraction amplitude of WT (*left*) and TEVs-TTN (*right*) cardiomyocytes, expressing or not GFP-TEVp (N=3). **(G)** Duration of contraction of WT (*left*) and TEVs-TTN (*right*) cardiomyocytes,, expressing or not GFP-TEVp (N=3).

In summary, we find that titin cleavage induces progressive disassembly of sarcomeric components in the absence of noticeable cell death. In addition, targeted cardiomyocytes retain measurable contractile activity.

### Invariant proliferation of cardiomyocytes containing cleaved titin

So far, our data demonstrate that full titin cleavage causes sarcomere disassembly without compromising the global viability of targeted cardiomyocytes. To assess whether titin-cleavage-induced sarcomere disassembly induces cardiomyocyte proliferation, we measured several markers of cell cycle (**Figures 5A)**. Specifically, we studied nuclear incorporation of 5-ethynyl-2′-deoxyuridine (EdU) (**Figures 5B, S3A**,**B**) and presence of Aurora B (**Figures 5C, S3A**,**B**) as markers of DNA synthesis and mitosis, respectively^33^. Quantification indicates that expression of GFP-TEVp does not alter EdU nuclear incorporation (**Figures 5D**), nor the proportion of Aurora B positive cardiomyocytes (**Figures 5E**). Accordingly, we observe that the fraction of binucleated AAV-infected TEVs-TTN cardiomyocytes is comparable to that of the controls (**Figure 5F**,**G**). Altogether, our results indicate that titin-induced sarcomere disassembly does not promote proliferation of neonatal cardiomyocytes *in vitro*.

**Figure 5.**
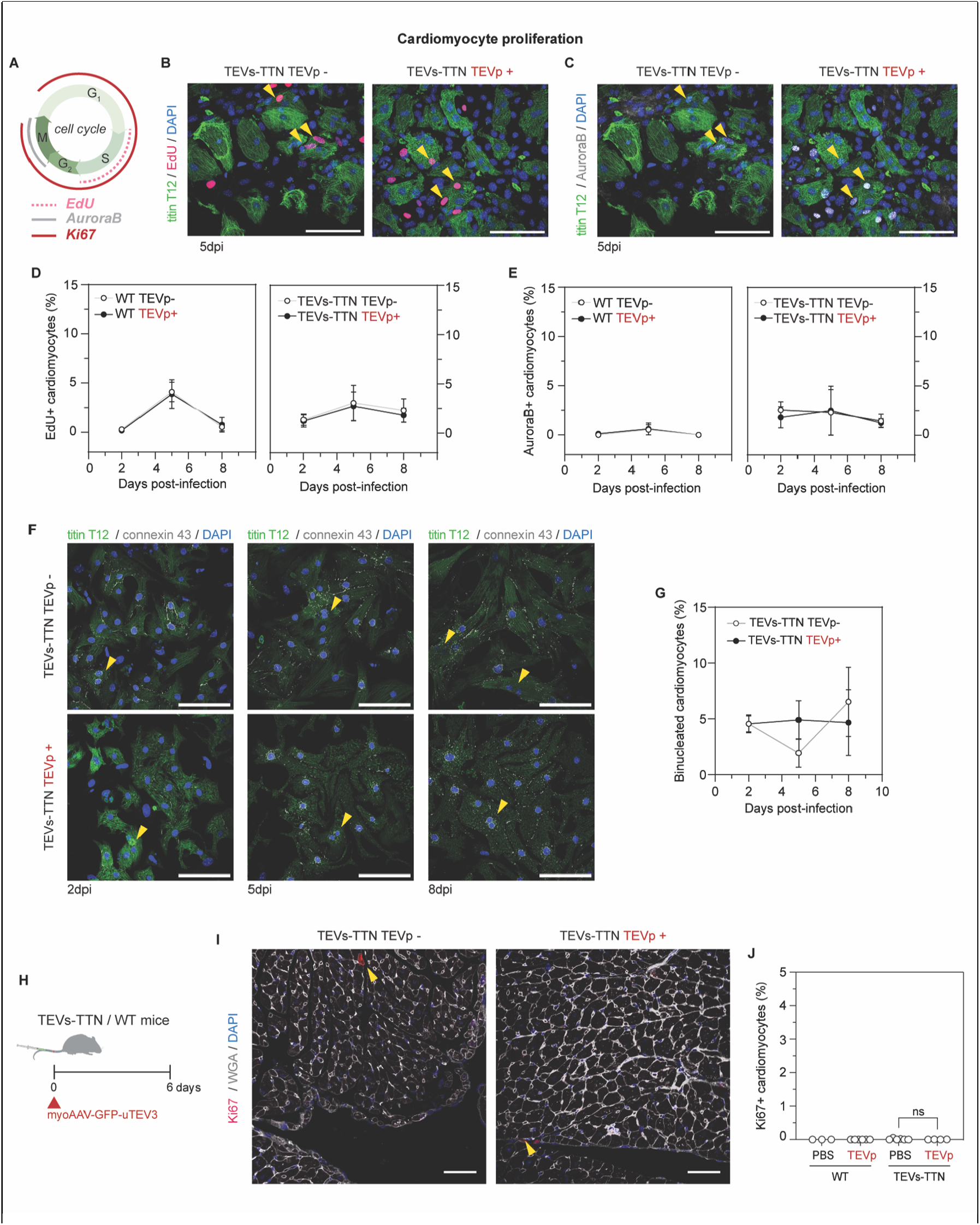
Examination of cardiomyocyte proliferation markers. **(A)** Schematic representation of markers of different stages of the cell cycle. **(B)** Representative images of EdU (*red*) incorporation in TEVs-TTN cardiomyocytes nuclei (yellow arrowheads) in samples infected or not with GFP-TEVp-expressing AAV6 vectors. Scale bars, 100 µm. **(C)** Representative images of Aurora B (*gray*)-positive nuclei in TEVs-TTN cardiomyocytes (yellow arrowheads) expressing or not with GFP-TEVp. Scale bars, 100 µm. **(D)** Quantification of EdU incorporation in WT (*left*) and TEVs-TTN (*right*) cardiomyocytes expressing or not GFP-TEVp. **(E)** Quantification of Aurora B-positive WT (*left*) and TEVs-TTN (*right*) cardiomyocytes expressing or not GFP-TEVp. **(F)** Representative images showing binucleated TEVs-TTN cardiomyocytes (yellow arrow heads). Scale bars, 100 µm. **(G)** Quantification of binucleated TEVs-TTN cardiomyocytes. **(H)** Schematic representation of titin-cleavage induction *in vivo*. **(I)** Representative immunodetection of Ki67 proliferation marker in ventricular myocardium of TEVs-TTN mice expressing or not GFP-TEVp. Scale bars, 50 µm. **(J)** Quantification of Ki67-positive cardiomyocytes in ventricular myocardium (N = 3-6).

We also investigated the effect of titin cleavage on adult cardiomyocyte proliferation by expressing GFP-TEVp in the myocardium of 9-19-week-old WT and TEVs-TTN mice (**Figure 5H**) under conditions that result in different levels of sarcomere disassembly in 30% of cardiomyocytes in TEVs-TTN mice^34^. As expected, proliferation analysis using Ki-67 as a cell cycle marker^35^ shows less than 0.05% proliferative adult cardiomyocytes in basal conditions; this fraction is not affected by expression of GFP-TEVp (**Figures 5I**,**J, S3C**). This result indicates that titin-cleavage-induced sarcomere disassembly does not trigger proliferation of adult cardiomyocytes *in vivo*, in agreement with our observations *in vitro*.

## Discussion

The cardiac muscle has limited capacity for renewal, which results in important functional deficits in the many sick hearts that experience loss of cardiomyocytes, regardless of the underlying etiology^36,37^. Opening new windows of opportunity to treat these highly prevalent heart diseases, in recent years several factors have been identified that can stimulate cardiomyocyte proliferation and promote cardiac regeneration, including inhibitors of cell signaling pathways, growth factors, and miRNAs ^13–18^. A common feature of cardiomyocyte division triggered by these factors, very much resembling natural cardiomyocyte proliferation, is the induction of a more dedifferentiated state of cardiomyocytes that includes disassembly of sarcomeres. Indeed, sarcomere disassembly itself has even been proposed to be a bona fide regulator of cell cycle progression^20^. Supporting this mechanistic role, overexpression of actin-capping adducin proteins induces sarcomere disassembly that, under some conditions, correlates with increased cardiomyocyte proliferative capacity^22^. Considering these results, we reasoned that the potential for cardiomyocyte regeneration could be also unlocked by sarcomere disassembly resulting from titin cleavage. Although our work does indeed demonstrate that titin cleavage causes disassembly of sarcomeres in the absence of noticeable cell stress or death, we find no evidence of accompanying proliferation of targeted neonatal cardiomyocytes *in vitro* or adult counterparts *in vivo*.

The mechanisms driving sarcomere disassembly during mammalian cardiomyocyte proliferation remain largely unknown although they seem to be adapted to the physiological state of cells (i.e. different mechanisms in, for example, embryonic *versus* neonatal cardiomyocytes)^20,22,38^. In our experiments, extensive titin cleavage is detected in GFP-TEVp-expressing TEVs-TTN neonatal cardiomyocytes as early as 2 dpi (**Figure 1F**,**G**). At this time, disassembly of the Z-disk into structures resembling Z-bodies is already evident, as shown by the location of *α*-actinin in immunofluorescence (**Figure 3**). Similarly, troponin T staining reveals disassembly of thin filaments in GFP-TEVp-expressing TEVs-TTN cardiomyocytes, although the distribution of the protein in targeted cells remains more fibrillar-like than for the other sarcomeric proteins we have studied (**Figures 2,3**). From 5 dpi, we find complete titin cleavage correlating with higher proportion of fully disassembled sarcomeres (**Figures 1F,G; 2D**). Intriguingly, T12 staining, which labels the Z-disk portion of titin, shows in targeted cells a non-sarcomeric pattern that is distinct from *α*-actinin’s (**Figures 2, 3**). This observation suggests that Z-disk protein complexes dissociate upon titin cleavage, and that the resulting protein components localize to different regions of the cell. A similar situation is observed for the titin fragments originating from GFP-TEVp activity (compare the diffuse M-band and spotted Z-disk stainings of titin in AAV-infected TEVs-TTN cardiomyocytes, **Figures 2B, 3C**). Remarkably, different location of titin regions is also detected during natural disassembly of sarcomeres in cycling embryonic cardiomyocytes ^39^. In this regard, it is hard to conceive how titin Z-disk and M-line titin epitopes, even if distant, can show different subcellular localization in cardiomyocytes undergoing metaphase without invoking proteolysis of titin^39^. In fact, several proteases targeting titin have been proposed to contribute to sarcomere disassembly^20^. Among them, MMP-2 has been identified as a fundamental mediator of oncostatin M-induced sarcomere disassembly and cardiomyocyte proliferation^21,40^. Remarkably, oncostatin M induces loss of striated pattern of both Z-disk and M-line regions of the sarcomere in a highly similar manner to the one we detect upon GFP-TEVp-mediated titin cleavage.

In combination our results indicate that titin-cleavage-induced sarcomere disassembly, despite not triggering cardiomyocyte proliferation, does recapitulate features that are also observed in proliferating cardiomyocytes. Whether proteolysis of sarcomeric components is required for sarcomere disassembly and cardiomyocyte proliferation remains controversial^21,38,39^. Arguably, extensive proteolysis would require rapid and energy-costing re-synthesis of sarcomeric components to limit the loss of contractile function of the myocardium. Our work adds a relevant counter-argument in this regard, since we demonstrate that just a single proteolytic cut in titin is enough to trigger sarcomere disassembly. Considering the well-known cardiac titin isoform switch that happens during late embryonic/early postnatal development^41^, it is tempting to speculate that titin cleavage during these proliferative stages could contribute to sarcomere disassembly of dividing cardiomyocytes while ensuring replacement of titin isoforms.

Importantly, our model of titin-cleavage-induced sarcomere disassembly also results in concomitant reduction of cardiomyocyte-cardiomyocyte adhesion strength and alteration of connexin 43 localization (**Figure 5F**, more extensive analysis *in vitro* and *in vivo* in^34^). Connexin 43 is the main component of gap junctions within intercalated discs, complex structures responsible for rapid and coordinated electromechanical coupling between neighboring cardiomyocytes^42^. Since cell-cell contacts appear to be preserved in naturally cycling cardiomyocytes^39^, the possibility exists that loss of cell adhesion upon titin cleavage somehow opposes cell proliferation in our model. Considering our results, it will be interesting to investigate why sarcomere disassembly in naturally proliferating cardiomyocytes does not compromise cell adhesion as we have detected here. A contributing factor may be that cardiomyocytes undergoing natural cell division are surrounded by several counterparts with intact sarcomeres unlike in our experiments.

In conclusion, our work demonstrates that cleaving titin is sufficient to induce sarcomere disassembly in living cardiomyocytes. However, the lack of ensuing cardiomyocyte proliferation suggests that sarcomere disassembly per se is not enough to trigger cell cycle entry, at least for the type of disassembly triggered by titin cleavage. Future work will investigate if concurrent cleavage of titin can boost cardiomyocyte proliferation induced by known mitogenic signals.

## Methods

### Animal experimentation

Animals were handled in accordance with European and Spanish guidelines for animal welfare and with the recommendations in the Guide for the Care and Use of Laboratory Animals of the National Institutes of Health. The protocol was approved by the Ethics Committee of Animal Experiments of our institutional committees (project numbers PROEX 042/18 and PROEX 107.8/23). Neonates were used in our experiments with no consideration of sex. Only male mice were used for *in vivo* experiments.

### Adeno associated virus production

Different TEVp constructs were expressed in adeno-associated virus (AAV) under the control of the CMV promoter. Specifically, AAV-compatible vectors CMV-mCherry-TEV and CMV-Tomato-TEV were from Addgene (#58868 and #58872, respectively). Plasmids CMV-GFP-uTEV3 and CMV-mCherry-uTEV3 encoding an improved TEVp variant were produced by us and can be obtained from Addgene (#234320 and #238000, respectively) ^31^. Production and quantification of AAV serotype 6 and AAV-MYO^43^ particles have been described elsewhere^28,34^. Titers are reported as viral genomes per milliliter (vg/mL).

### Cardiomyocyte isolation and infection

Neonatal mouse cardiomyocytes were isolated from WT and TEVs-TTN mice. Dissected hearts from neonatal (P1-3) were minced with a scissor in cold Hanks’ Balanced Salt Solution (HBSS) and dissociated using the Pierce Primary Cardiomyocyte Isolation Kit (Thermo Scientific^TM^Pierce^TM^, 88281) in 0.21 ml enzyme mix/heart in a 2 ml reaction tube for 20 minutes at 37°C with occasional gentle shaking. The pellet was centrifuged, washed twice with HBSS and resuspended in 0.5 ml cardiomyocyte medium (DMEM for Primary Cell Isolation containing 10% heat-inactivated FBS and 1% penicillin/streptomycin) per heart and plated onto MatTek dishes covered with matrigel solution (Fisher Scientific, 11553620). Cardiomyocytes were infected overnight with AAV6-expressing TEVp constructs 24-48 hours after plating.

### Cardiomyocyte live imaging

Isolated cardiomyocytes seeded in p35 culture plates (Mattek, P35G-0-10-C) were used for live imaging. After 2, 5, 8, and 12 dpis the culture media were replaced by DMEM without phenol red (Gibco™, 31053028) containing 10% heat-inactivated FBS and 1% penicillin/streptomycin. Time-lapse imaging was performed at 37°C and 5% CO_2_ using a Plan Apo λ 20x/0.75 objective in a Nikon ECLIPSE Ti epifluorescence microscope coupled to an Orca ER hamamatsu CCD camera. GFP-TEVp expression was quantified by capturing native green fluorescence.

### Quantification of titin cleavage

2×10^6^ cardiomyocytes were seeded in p35 culture plates. Cell lysates were obtained in 150 *μ*l of sample buffer (3% SDS, 50 mM TrisCl pH 6.8) supplemented with 50 mM N-ethylmaleimide (Sigma-Aldrich, E3876) and protease inhibitor Cocktail Set III (Millipore). Samples were diluted in 0.06% bromophenol blue and 10% glycerol and run on 1.8% polyacrylamide SDS-PAGE gels. Titin proteins were visualized by SYPRO Ruby staining (ThermoFisher, S21900) and relative band intensities were measured using QuantityOne Software.

### Atomic force microscopy

AFM measurements were performed with a commercial JPK NanoWizard III instrument (Bruker-JPK, Berlin, Germany) coupled to an inverted Axio Observer A1 optical microscope (Carl Zeiss, Germany). Cardiomyocytes were permeabilized with 0.2% (v/v) Triton X-100 in 1% (w/v) BSA for 15 minutes and were probed at room temperature in PBS using rectangular Si_3_N_4_ AFM cantilevers with silicon tips of radius < 15 nm and 0.1 N m^-1^ nominal spring constant (BioLever mini, Bruker). Cantilevers were calibrated by the thermal fluctuation method^44,45^. Force *versus* distance curves were recorded in contact mode on the surface of single cardiomyocytes to determine Young’s moduli. For each cell, approximately 200 force-distance curves were recorded. Approach-retract cycles were performed at 10 µm s^-1^ (5 µm ramp size and 0.5 nN setpoint force). Young’s modulus values (*E*) were calculated using the JPK Data Analysis software, by fitting the approach section of the curve to the Hertz model for a paraboloid using Equation 1^46^:

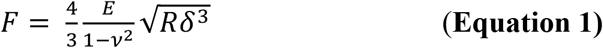

In Equation 1, *F* is the applied force, *R* is the probe radius, *ν* is Poisson’s ratio, and *δ* is the indentation. Cells mostly consist of water and are considered incompressible, hence a Poisson ratio of 0.5 was used.

### Immunocytochemistry and immunohistochemistry

Neonatal cardiomyocytes were fixed with 4% (v/v) paraformaldehyde (PFA) at room temperature (RT) for 10 min, permeabilized with 0.2% (v/v) Triton X-100 (Sigma-Aldrich) in 1% (w/v) BSA (Sigma-Aldrich) for 5 min and blocked in SB buffer (PBS containing 10% FBS and 1.5% BSA) for 1 hour at RT. Following incubation with primary antibodies at 4°C overnight, samples were incubated with secondary antibodies and 5 *μ*g/mL 4′,6-diamidino-2-phenylindole dihydrochloride (DAPI) to mark the location of the nuclei at RT for 1 hour (antibodies are reported in the **Table S1**). Fluoromont G (bioNova, 0100-01) was used as mounting medium and allowed to cure overnight. Images were taken using confocal microscope Zeiss LSM700 with Plan-Apochromat 40x/1.3 Oil DIC M27. Regarding myocardial samples, ventricles were fixed in PFA overnight at 4ºC. After washing in PBS, the tissue was subsequently treated with 10% and 30% sucrose, embedded in optimum cutting temperature (O.C.T.) (Tissue-Tek® Compound 4583, Sakura) and sectioned at 8 μm with a cryostat (Leica). Transverse sections of the middle portion of each segment were permeabilized with 0.3% Triton X-100 in PBS and blocked in SB buffer. Sections were first incubated with anti Ki67 at 4°C overnight and after washing, they were incubated with the anti-rat IgG -Alexa Fluor^®^ 555 Plus and wheat germ agglutinin (WGA) at RT for 1.5 hour (**Table S1**). After the secondary antibody incubation, sections were stained with DAPI. Mowiol 4-88 (Sigma-Aldrich) was used as mounting medium and allowed to cure overnight. Images were taken using confocal microscope Nikon A1R with a Plan Apo 40x/1.3 Oil objective.

### Image analysis

We used Fiji platform^47^ and custom script plugins. To measure GFP-TEVp levels, cardiomyocytes were identified using sarcomere markers and were segmented manually. GFP-positive cells were counted, and regions containing green fluorescent cardiomyocytes were selected to measure GFP intensity. Sarcomere organization was estimated measuring anisotropy of T12 signal using FibrilTool^48^. Only cardiomyocytes showing T12 staining were used in the analysis of cell cycle markers *in vitro*. Binucleated cardiomyocytes were quantified based on the presence of double DAPI positive nuclei in manually segmented cells, using connexin 43 as a membrane marker. For TEVs-TTN-TEVp, which exhibit a reduction in connexin 43, cell segmentation is facilitated because cells have less intimate contacts with their neighbors^34^. Ki67-positive cardiomyocytes in the myocardium were quantified using a custom script plugin building on the Cellpose plugin^49^ for general cellular segmentation based on the WGA membrane marker, and the Stardist plugin^50^ for nuclear segmentation. Cellular area larger than 50 µm^2^ was used to quantify positive cardiomyocytes.

### Cardiomyocytes metabolic activity

Cardiomyocyte viability was examined using the 3-[4,5-dimethylthiazol-2-yl]-2,5-diphenyl tetrazolium bromide (MTT) assay (Roche, 11465007001). Briefly, neonatal cardiomyocytes were seeded on 96-well plates (Mattek, P96G-1.5-5-F). At 5 dpi, cell medium was replaced by DMEM without phenol red (Gibco™, 31053028) and 0.5 mg/ml MTT was added. Cells were incubated at 37 ºC and 5% CO2 for 4 hours. Next, solubilization buffer (10% SDS in 0.01 M HCl) was added to the cells and used to solubilize formazan crystals. Absorbance was determined using a spectrophotometer (SPECTROstar Nano Microplate Reader, BMGLABTECH Germany) at 570 nm and background absorbance at 690 nm was subtracted. Final absorbance signal is proportional to the number of metabolically active cells. Cells exposed to 1 mM hydrogen peroxide (H_2_O_2_) were taken as control, non-viable samples.

### TUNEL assay in neonatal cardiomyocytes

Apoptotic damage was examined using a TUNEL assay kit following the recommendations of the provider (Roche, 11684795910). Briefly, neonatal cardiomyocytes were cultured on 24-well plates (Mattek, P24G-1.5-13-F). At 5 dpi, the cells were fixed with PFA for 10 min, permeabilized with 0.2% (v/v) Triton X-100 in 1% (w/v) BSA for 5 min. Treatment with 1 ng/ml DNase I before incubation with enzyme solution (TdT) served as positive control. As negative control, samples incubated with Label Solution (fluorescein-dUTP) alone wwere used. Fluorescent images were acquired using a Zeiss LSM700 confocal microscope with Plan-Apochromat 40x/1.3 Oil DIC M27 objective.

### Beating analysis using MYOCYTER

Short videos of spontaneously beating single cardiomyocytes, lasting 14.26 seconds at 33.3 frames per second, were analyzed from 10-20 beating cardiomyocytes. MYOCYTER, an open-source macro for ImageJ, was used to determine contraction parameters ^51^. The macro measures contraction from the image differences between the resting state (reference image) and the subsequent frames. Peak times were estimated at 20% amplitude.

### EdU assay

Nuclear 5-ethynyl-2′-deoxyuridine (EdU) incorporation was used to evaluate cellular proliferation. EdU assays was performed with the Click-iT™ Plus EdU Alexa Fluor™ 647 Imaging Kit according to the manufacturer’s protocol (ThermoFisher). EdU was added to cardiomyocyte cultures 3 hours before cell fixation according to the manufacturer’s protocol.

## Statistical analysis

In all figures, measurements are reported as mean ± standard error of the mean (SEM). The number of independent experiments is specified in the figure legends. In the case of *in vitro* experiments, N refers to the number of independent isolations of cardiomyocytes. Statistical significance was evaluated using GraphPad Prism 10. Unpaired Student’s two-sided t-test were used to assess statistically significant differences between groups. Differences were considered statistically significant at p < 0.05. Non-significant differences are indicated as “ns”.

## Supporting information

Supplementary Movie S1 - TEVs-TTN cardiomyocytes 5 dpi

Supplementary Movie S2 - TEVs-TTN cardiomyocytes TEVp 5 dpi

## Authors contributions

MRP, MALU, RSR, EHG, JAC conceived and designed the experiments. MRP, MALU, NV, CML, CHL, WPF, ACD, FME executed and/or analyzed experiments. MIS and RG contributed materials and analysis tools. MRP and JAC drafted the manuscript. All authors revised and approved the final manuscript.

## Acknowledgments

We acknowledge the personnel from CNIC animal facility and viral vectors. Light microscopy was conducted at the CNIC Microscopy & Dynamic Imaging Unit. We thank Verónica Labrador-Cantarero for her support with imaging analysis. We acknowledge the feedback from many colleagues at CNIC, especially Florian Weinberger, José Luis de la Pompa, Silvia Martin-Puig, Hesham Sadek, and Miguel Torres and his team. We thank and Mauro Giacca for insightful feedback. We thank Dieter Fürst for providing the T12 antibody. We thank all members of the Molecular Mechanics of the Cardiovascular System team for their support and input. We thank Alba Pobes-Lagartos for excellent technical assistance.

## Data availability

The data relating to this article are available in the article itself or in its Supplementary material online. The beating analysis data are available upon reasonable request to the corresponding authors.

