## Supplementary Information for "Titin cleavage in living cardiomyocytes induces sarcomere disassembly but does not trigger cell proliferation"

[1]Centro Nacional de Investigaciones Cardiovasculares (CNIC), Calle de Melchor Fernández Almagro 3, 28029, Madrid, Spain.
[2]Louvain Institute of Biomolecular Science and Technology, Universite catholique de Louvain, Louvain-la-Neuve, Belgium.
[3]Instituto de Ciencia de Materiales de Madrid (ICMM), Madrid, Spain.
[4]University of Cambridge, Department of Chemistry, Lensfield Road, Lensfield Road CB2 1EW, Cambridge, UK.

**This file includes:**

**- Supplementary Figures S1-S3**

**- Supplementary Table S1**

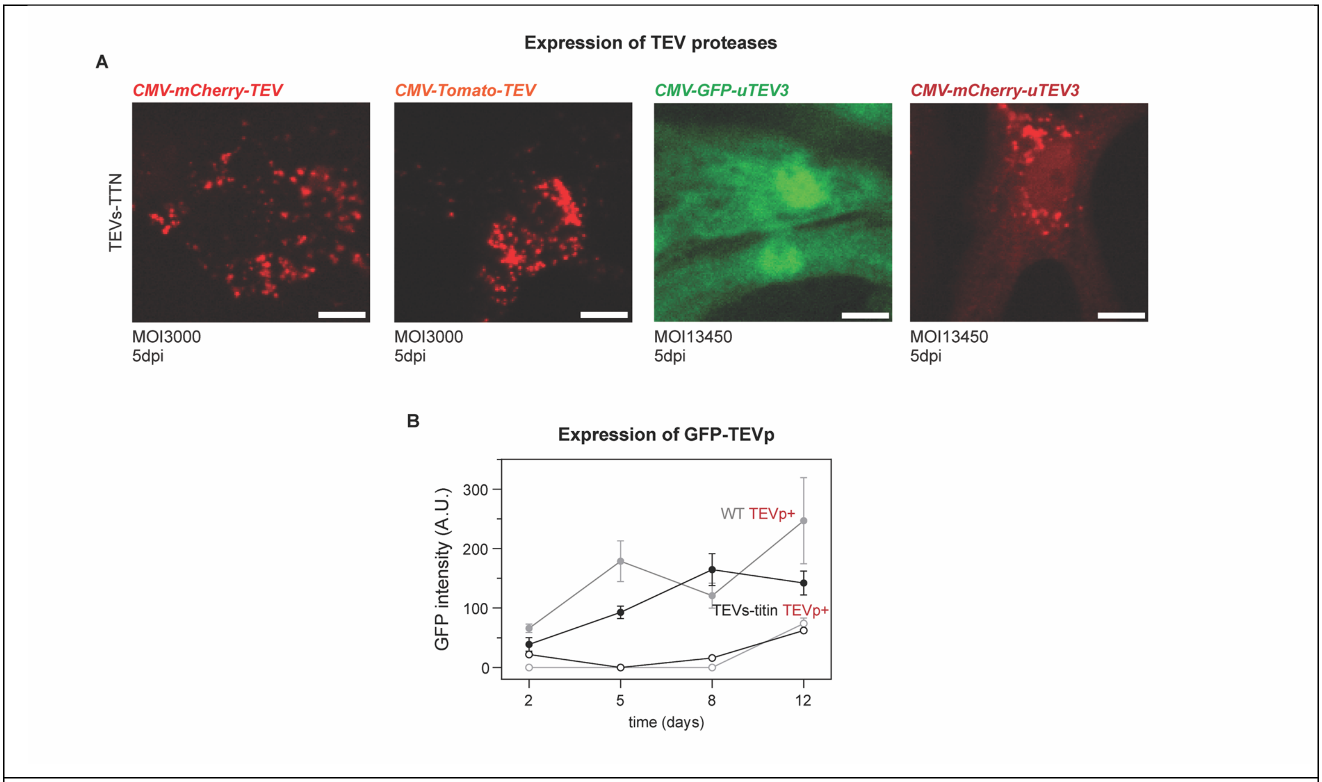


**Figure S1. Expression of GFP-TEVp in neonatal cardiomyocytes. (A)** Representative images of the screening of expression of different Tobacco Etch Virus (TEV) proteases in neonatal cardiomyocytes. Multiplicity of infection (MOI) used in the experiments is indicated. Scale bars: 10 µm. **(B)** GFP-uTEV3 (referred to in the paper as GFP-TEVp) expression count in epifluorescence images of living cardiomyocytes. Each data point represents the mean of fluorescence intensity in positive cardiomyocytes at the different experimental times.

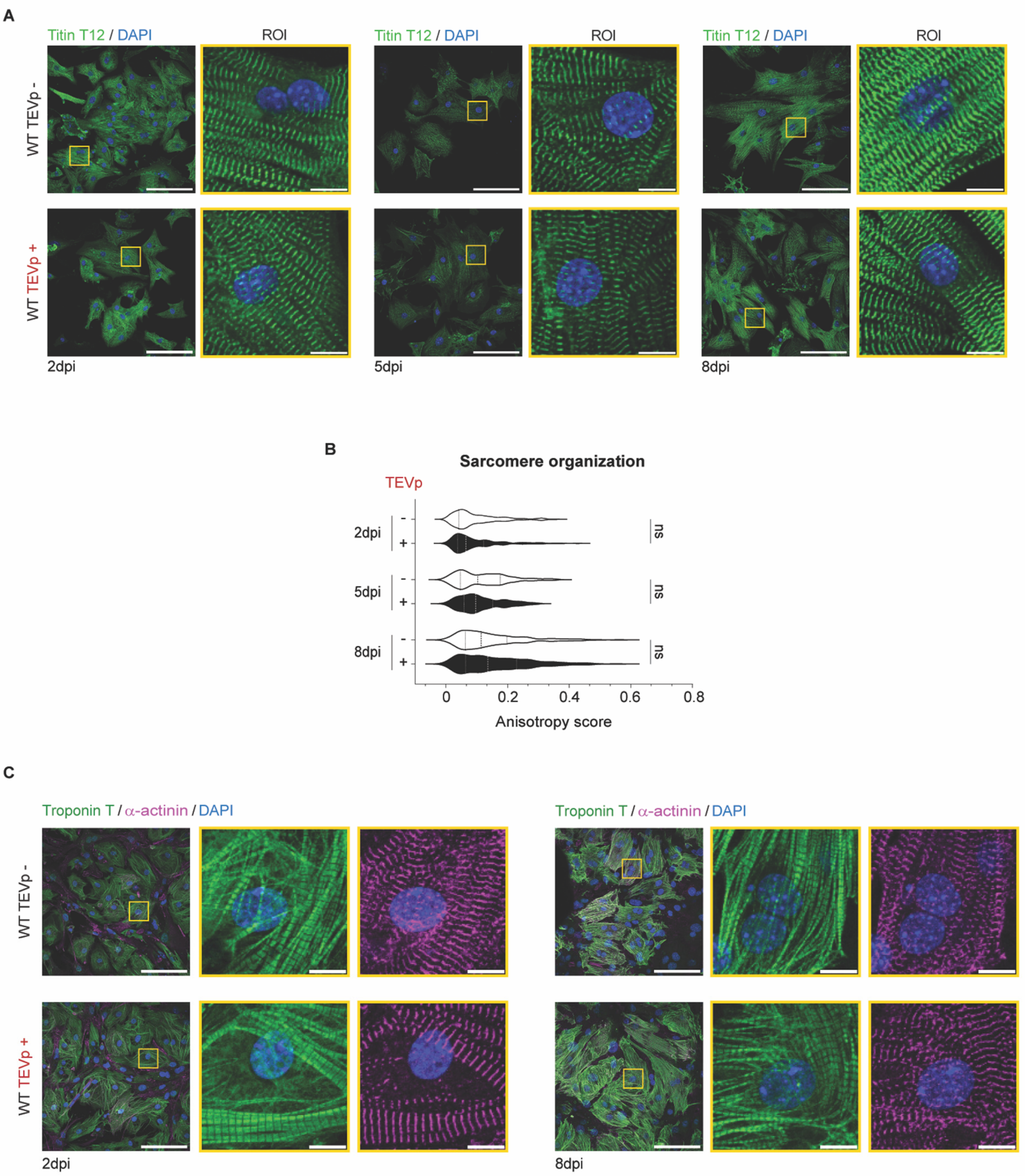


**Figure S2. Sarcomere structure upon GFP-TEVp expression in WT cardiomyocytes**. **(A)** Representative immunofluorescence images of WT cardiomyocytes expressing or not GFP-TEVp to monitor T12 localization. **(B)** Sarcomere organization quantified as anisotropy of T12 signal in immunofluorescences in WT cardiomyocytes expressing or not GFP-TEVp (N=2-3). **(C)** Representative immunofluorescence images of WT cardiomyocytes expressing or not GFP-TEVp focusing on thin filament and Z-disk structures. Scale bars, 100 µm. ROI scale bars, 10 µm.

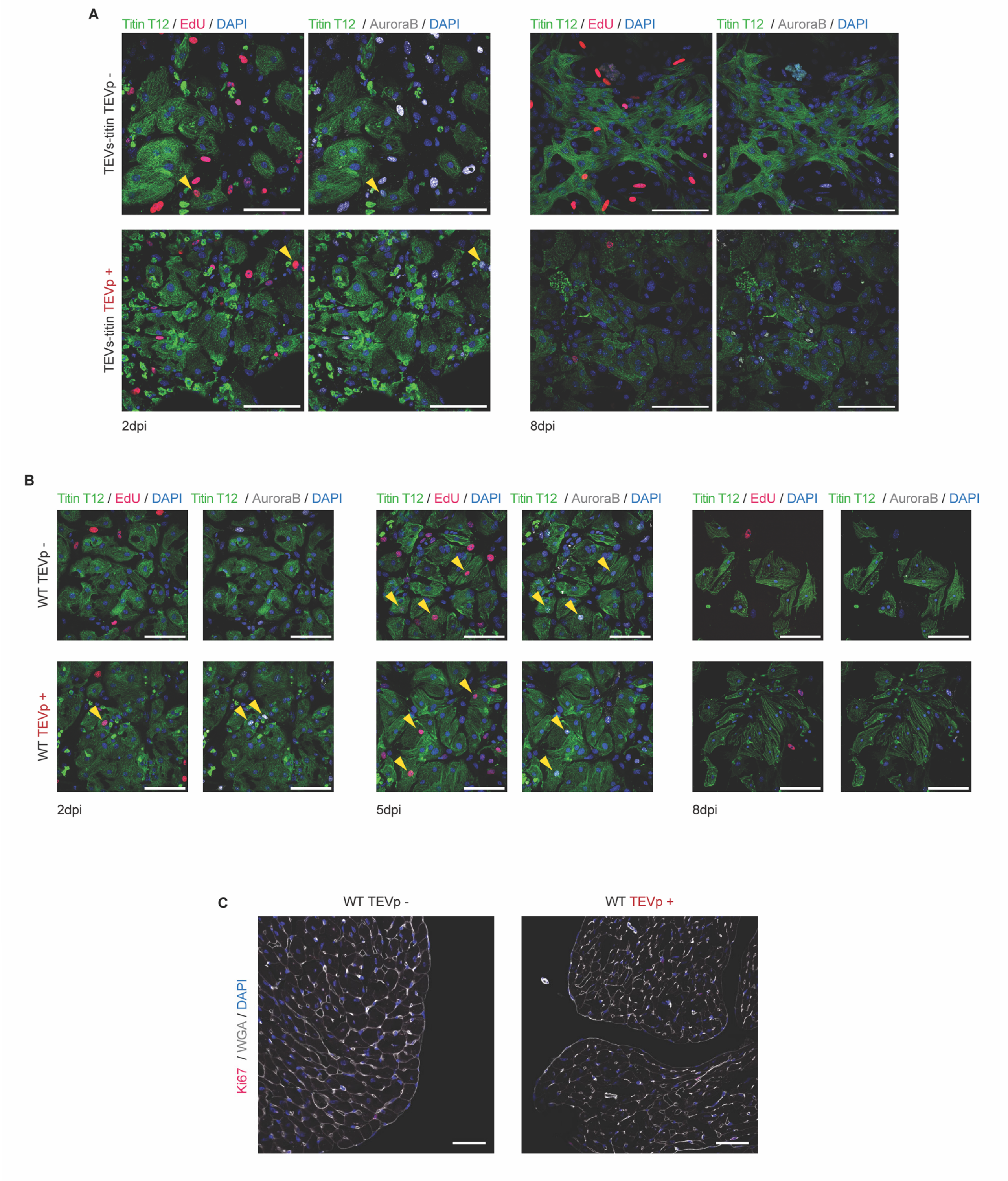


**Figure S3. Cardiomyocyte proliferation. (A)** Representative images of EdU incorporation and Aurora B positive cardiomyocytes (yellow arrowheads) in TEVs-TTN cells expressing or not GFP-TEVp. Scale bars, 100 µm. **(B)** Representative images of EdU incorporation and Aurora B positive cardiomyocytes (yellow arrowheads) in WT cells expressing or not GFP-TEVp. Scale bars, 100 µm. **(C)** Immunodetection of ki67 in ventricular myocardium of WT mice expressing or not GFP-TEVp. Scale bars, 50 µm.

**Table S1. Staining reagents used for immunofluorescence**

| Reagent | Reference | Dilution (ICC) | Dilution (IHC) |
|---|---|---|---|
| **anti cleaved TEV site** | *Novus Biologicals* (NBP2-37831) | 1:100 | - |
| **anti titin T12 (Z-disc)** | provided by D.O. Fürst, Bonn[2] | 1:100 | - |
| **anti troponin T-Alexa Fluor® 488** | *Bioss Antibodies* (BS-10648R-A488) | 1:200 | - |
| **anti α-actinin** | *Sigma-Aldrich (*A7732) | 1:100 | - |
| **anti Titin TTN-9 (M-band)** | *Myomedix (TTN-9-y)* | 1:100 | - |
| **anti Aurora B** | *Abcam* (ab2254) | 1:100 | - |
| **anti Ki67-PE** | *ThermoFisher Scientific* (12-5698-82) | - | 1:100 |
| **wheat germ agglutinin (WGA) –Alexa Fluor® 488** | *ThermoFisher Scientific* (W11261) | - | 1:250 |
| **DAPI** | *Sigma-Aldrich* (MBD0015) | 5 µg/mL | 0.5 µg/ml |
| **anti rabbit IgG-Alexa Fluor® 647** | *ThermoFisher Scientific* (A-21245) | 1:500 | - |
| **anti rabbit IgG-Alexa Fluor® 488** | *ThermoFisher Scientific* (A-21441) | 1:500 | - |
| **anti rabbit IgG-Alexa Fluor® 555** | *ThermoFisher Scientific* (A48263) | 1:500 | - |
| **anti mouse IgG-Alexa Fluor® 647** | *ThermoFisher Scientific* (A-21463) | 1:500 | - |
| **anti mouse IgG-Alexa Fluor® 488** | *ThermoFisher Scientific* (A-11029) | 1:500 | - |
| **anti chicken IgY-Alexa Fluor® 647** | *Abcam* (ab150171) | 1:500 | - |
| **anti rat IgG-Alexa Fluor® 555 Plus** | *ThermoFisher Scientific* (A48263) | - | 1:500 |

ICC=Immunocytochemistry

IHC=Immunohistochemistry
